# *In silico* validation and fabrication of matrix diffusion based polymeric transdermal films for repurposing gabapentin hydrochloride in oncogenic neuropathic pain

**DOI:** 10.1101/2020.12.01.406041

**Authors:** Manisha Singh, Shriya Agarwal, Pranav Pancham, Harleen Kaur, Vinayak Agarwal, Ramneek Kaur, Shalini Mani

## Abstract

**Background:** Gabapentin (GBP) is an FDA approved drug for the treatment of partial and secondary generalized seizures, apart from also being used for diabetic neuropathy. GBP displays highly intricate mechanism of action and its inhibitory response in elevated antagonism of NMDA (N-methyl-D-aspartate receptor) receptor and has potential in controlling neuropathic pain of cancer origin.

**Objective:** Therefore, in the present study, we have selected BCATc (Pyridoxal 5’-phosphate dependent branched-chain aminotransferase cytosolic) enzyme that is highly expressed in neuropathic stress conditions and have analysed the GBP as its competitive inhibitor by modeling, docking and checking its pharmacokinetic suitability through ADMET. Though in this study the results exhibited higher efficacy of GBP in controlling neuropathic pain, the drug shows certain potential therapeutic limitations like shorter half-life, repetitive dosing, high inter subjective variability.

**Methods:** Therefore, a suitable and equally efficacious drug delivery method was also designed and developed by loading GBP transdermal patches (GBP-TDP) by solvent evaporation method using PVP and HPMC in ratio of 2:1 as a polymer base for reservoir type of TDP. Also, PEG 400 was used as a plasticizer and PVA (4%) was taken for backing membrane preparation and then the optimized GBP-TDP was subjected for physical characterization, optimization and *ex vivo* release kinetics.

**Results and conclusion:** The results showed desired specifications with uneven and flaky surface appearance giving avenue for controlled release of the drugs with 75.58% of drug release in 12 hrs., further suggesting that GBP-TDP can be used as an effective tool against diabetic neuropathy pain.

## 1. INTRODUCTION

The oncological pain sensation arising as a direct consequence of neuronal damage or affecting the somatosensory cortex, either at peripheral (PNS) or central nervous system (CNS) is marked as neuropathic pain and is represented by patients as a debilitating pain apart from being the unrelenting type. This type of excruciating pain compared as equivalent to peripheral nerve injuries is noticed as events in the primary afferents[1], caused by peripheral sensitization including decreased threshold (allodynia; pain in response to innocuous stimulus), increased response to supra-threshold (hyperalgesia) and spontaneous activity of nociceptors[2]. Subsequently, a series of molecular events in the CNS is associated with central sensitization resulting in synaptic transmission altercation, increased excitatory synaptic processes and attenuation in inhibitory processes. In such kind of pain existence, the recommended first line treatments for neuropathic pain are anticonvulsants, antidepressants and it’s been observed that usually non-steroidal anti-inflammatory drugs (NSAIDs) do not help much in providing sustained relief from pain. Further, it’s been also reported that serotine-noradrenaline uptake inhibitors like GBP and duloxetine provide instant and effective relief in shortest duration followed by Tramadol, opioid analgesics (second line treatment), the most recent one-cannabinoids (third line treatment) and lastly, lidocaine, capsaicin, carbamazepine and ziconotide as a fourth-line of treatment[3, 4]. Similarly, in the series it’s been seen that Gabapentin (GBP), an FDA approved second-generation anti-epileptic drug has demonstrated the efficacy in treating various types of neuropathic pain including painful diabetic neuropathy, trigeminal neuralgia and bipolar disorders. The molecular mechanism(s) by which GBP, a structural analog of GABA (Gamma Aminobutyric Acid), exerts its anticonvulsant action is unknown but it has been reported that the pathway mechanism involves direct inhibition of α2δ subunit of a voltage dependent calcium channel resulting in decline of presynaptic Ca^2+^ influx and subsequent, release of excitatory neurotransmitters like glutamate[5]. GBP is found to be involved in inhibiting ectopic discharge activity from damaged peripheral nerves and these distinct aspects, sequentially suggests collective analgesic expression of GBP in chronic neuropathic pain. Moreover, GBP was originally illustrated as GABA mimetic to accomplish a primary purpose of crossing blood brain barrier (BBB) and interaction with GABAergic systems to elevate GABA mediated inhibitions[6]. Amidst various theories; effects on CNS due to either elevated antagonism of NMDA (N-methyl-D-aspartate receptor) receptor, inhibitory response of GABA–mediated pathways and antagonism of existing calcium channels in CNS or inhibition of peripheral neurons is considered as one of the most widely accepted expression of working by GBP[7]. Further, it has been noticed that NMDA receptor complex is classed as a ligand – gated ion channels, responsible for regulating Ca^2+^ ion influx when activated[8]. This complex expresses several active sites for specific binding with different ligand such as proton – sensitive site, redox modulatory site, phencyclidine binding site and strychnine-insensitive glycine binding site. Also, one of the primary subunit of NMDA receptor complex is the α_2_δ, a calcium ion channel mediated voltage – dependent complex that associates itself with GBP to elicit anti-hyperalgesic effects[9]. This activity results in decline of Ca^2+^ influx that guides the release of excitatory amino acids further, leading to decreased AMPA (α-Amino-3-hydroxy-5-methyl-4-isoxazolepropionic acid) receptor activation and conventional release of noradrenaline in the brain. Similarly, BCATc (Pyridoxal 5’-phosphate dependent branched chain aminotransferase cytosolic) gene has been reported to exhibit a profound expression and regulation in neuropathic stress condition[10, 11] Now, the metabolic theory of GBP action is through the inhibition of branched chain amino acid transamination and was also demonstrated by Hutson and coworkers that GBP is a competitive inhibitor of hBCAT_C_ with a K_i_ (inhibition constant) similar to the K_m_ (Michaelis constant) of Leucine[12]. But after reviewing the literature we figured out that there has been a lack of structural information about the specificity of GBP to hBCATc. Therefore, in the present study, we have complexed GBP with hBCATc and provided the understanding for the molecular interactions for the specificity of GBP with hBCATc. Furthermore, since we also noticed that pharmacokinetically the effectiveness of GBP in neuropathic pain gets decreased due to its dose dependent oral bioavailability that exist as an average of about 60% at a 300 mg dose to about 35% with dose increase[13] It also exhibits larger inter subject variability (27-66%) in plasma, rapid clearance of the drug along with shorter plasma half-life (5-7 hrs.) with multiple dosing leading to the increased in patients non-compliance apart from having multiple systemic side effects like - somnolence, asthenia, ataxia and dizziness[14]. Therefore, to have an effective and targeted delivery with retained structural specificity of GBP interaction to optimize efficacy and reduce its side effects was needed. Hence, to overcome these limitations we have developed transdermal patches (TDP) of GBP (GBP-TDP) that offers a non-invasive, and an alternative route of administration for rapid GBP delivery to the targeted site through directly getting in to systemic circulation and circumventing the liver metabolism. Besides, being thermodynamically stable, lower pore size and a large interfacial area GBP-TDP makes an appropriate drug delivery tool to enhance uptake across epithelial layer of skin and eventually increasing the bioavailability.

## 2. MATERIALS AND METHODOLOGY

### 2.1 *In silico* validation by targeting hBCATc in the oncogenic neuropathic pains

As discussed above, BCATs has pyridoxal 5’-phosphate (PLP) as a cofactor that catalyzes the reversible transamination of the short aliphatic branched-chain amino acids to their respective α-keto acids[15]. Many studies reported that BCAT isoenzymes further causes BCAT-dependent nitrogen shuttle catalysis and the conversion of astrocyte citric acid cycle intermediated to glutamate. Therefore, by the inhibition of hBCATc, found exclusively in brain, slows down the synthesis and reduces the amount of glutamate released during excitation in the neuronal tissues[16]. This effect can be useful for treatment of a wide variety of neurological driven disorders involving disturbances of the glutamatergic system[17]. Although there are many docking studies reported on aminotransferase inhibitors, but to our knowledge there are no clear evidences reported for analyses of toxicity of anticonvulsant and non metabolizable leucine analog of GBP. Our approach was to develop a three-dimensional homology model of the target hBCATc and to analyse the activity of the inhibitor GBP against it and later checking the predicted toxicity and pharmacokinetics parameters of the inhibitor by ADME data set.

#### 2.1.1 Homology modeling

Homology modeling of proteins offers immense value for interpretation of the relationships between the sequences, there structures and functions[18]. In this study, query sequence or the target protein (hBCATc) was aligned with the known template structure and later modeled.The protein sequence was retrieved from UniProt (https://www.uniprot.org/) with ID: P54687 and then subjected to BLAST (Basic Local Alignment Search Tool) (https://blast.ncbi.nlm.nih.gov/Blast.cgi) through protein BLAST tool (BLASTp) against the non-redundant database to identify that protein with maximum query coverage and highest sequence similarity will be considered as the template for further steps of modeling[19]. Using MODELLER 9.24 version the template was aligned with the query, later modeled using python based application (version 3.7) and the best model was selected on the basis of DOPE score and GA341 score. Subsequently, the modeled protein was visualized through the visualization software UCSF Chimera 1.14. and then the best model was further validated by generating the Ramachandran Plot using the PROCHECK webserver that exhibits the data for the most favoured and non-favoured residues of the modeled protein[20] and then further taken for its binding site analysis. The inhibitors were also selected for this modeled target protein.

#### 2.1.2 *In silico* Molecular Docking

##### 2.1.2.1 Preparation of ligand (GBP) and modeled protein (2COJ)

The refinement of modeled protein (receptor) is an essential step towards achieving quality docking results and the 3D-X-ray diffracted crystal structure of 2COJ was obtained through modeling studies. Data from homology modeling suggested the best designed model of the hBCATc protein has the highest sequence similarity of 99.74%, was further taken as target for docking studies. Procured file was then cleansed by removing the water molecules, heat atoms, excising alternating conformations, inserting Kollman charges and adding polar hydrogen to the 3-D protein moiety using Python Molecular Viewer-1.5.6 and then the .pdb file was converted into .pdbqt file for grid formation. Additionally, the atomic co-ordinate of the ligand was retrieved from the database of the natural compounds PubChem (https://pubchem.ncbi.nlm.nih.gov) [21]. In our study, ligand (Gabapentin (2- [1- (aminomethyl) cyclohexyl] acetic acid)) having chemical formula: C_9_H_17_NO_2_ and PubChem CID: 3446 (https://pubchem.ncbi.nlm.nih.gov/compound/3446) was taken as an inhibitor of hBCATc protein. Ligand structure was processed via LigPrep Tool in Maestro 2015 and was neutralized at *p*H 7.0 ± 2.0. Later the moiety was employed to OPLS2005 force field algorithm to minimize the energy of the 3D structure. The target for docking studies against the ligand (GBP) was already modeled and was later optimized using the steepest descent method from the Gromacs package version 2020.3[22]

##### 2.1.2.2 Grid generation and active site molecular docking

The grid generation of the modeled protein structure is an explicit demarcation of a receptor region where active binding interaction can occur and this was performed by using AutoDockTools-1.5.6, after preparing the co-ordinate files for ligand and target. The interactions between the target and the ligand was analyzed using the Lamarckian Genetic Algorithm (LGA) and before proceeding for the docking, identifying the potential active binding sites on the target and other associated data related to interacting residues was performed using a webserver CASTp 3.0 (http://sts.bioe.uic.edu/castp/index.html?3trg) [23]. The grid maps defining the search regions on the target were calculated with Autogrid command and had the grid dimensions of 60 Å × 50 Å × 40 Å with a spacing of 0.669 Å in between the grid points. The optimized ligand now fits into the target site at the docking run that evaluated the binding energy in terms of kcal/mol using the hydrogen bonding, van der Waals, electrostatic interactions. The docked confirmations were then visualized using UCSF Chimera 1.14 and the best was estimated from the resultant log file of docking studies exhibiting the binding affinity of all the docked conformations [24].

#### 2.1.3 Pharmacokinetic study through ADMET

The pharmacokinetic parameters (Absorption, Distribution, Metabolism, Excretion and Toxicity) predicted details helps in understanding the drug concentration in blood/plasma along with its bioavailability, metabolism and toxicity. A balanced and stable predictive model for ADMET parameters depends ideally on selection of the right mathematical approach, correct molecular descriptors for particular ADMET endpoints[25]. Therefore, in this study, ADMET analysis was carried out to check the pharmacokinetic behaviour of the ligand when it enters the body tissues and this was completed using the SwissADME (http://www.swissadme.ch) web tool with ADMET model of Discovery Studio software to access the drug-like properties of the ligand used in the docking studies[26].

### 2.2 Preparation of Transdermal patch (TDP)

GBP was a gift from Sun Pharmaceuticals Ltd. Polyvinyl alcohol (PVA), polyvinyl pyrolidone (PVP), Ecopol L100 and S100, (Hydroxypropyl) methyl cellulose (HPMC), Carbopol, Dibutyl phthalate (DBP), and polyethylene glycol 400 (PEG 400) were purchased from Sigma Aldrich Inc. Franz Diffusion Cell was purchased from Rama scientific. All other chemicals were of analytical grade.

#### 2.2.1 Quantitative analysis of GBP

Quantitative analysis of GBP was performed by using reversed-phase, high-performance liquid chromatographic (Waters) method. The chromatographic conditions were: column C_18_ with detection at 210 nm; temperature was fixed at 35°C. A stock solution of GBP (10 mg/mL) was prepared in mobile phase, with subsequent dilutions (2, 4, 6, 8, 10 mg/mL). The isocratic mobile phase consisted of methanol, acetonitrile and 0.028 M potassium dihydrogen phosphate (25:10:65), which was degassed by passing through a 0.45 μm Millipore filter and sonicated for 10 min and pumped at a flow rate of 1.0 ml/ min[27].

#### 2.2.2 Preparation of backing membrane and transdermal patches

Backing membrane was prepared by dissolving polyvinyl alcohol (PVA) in different concentrations of distilled water to form 2% and 4% (w/v) solutions and were kept on stirring for 6 hrs. to get homogeneous solution. After that, prepared solutions were poured on anumbra petri dishes and left for drying at 45°C for 24 hrs.[28].

To prepare the TDP, different combinations of different polymers (Ecopol L1100 and S100, Polyvinyl pyrolidone (PVP), Hydroxypropyl methyl cellulose (HPMC) were taken and dissolved in different ratios of suitable solvents (water and ethanol). After optimizing the various concentrations of polymers used, 300 mg of GBP was added in the polymer solution. Moreover, the plasticizer (PEG 400 or DBP) was also added (200 μL) in the formulation to enhance the flexibility and processability of the same followed by the addition of DMSO (200 μL) as a permeation enhancer while stirring. The presence of plasticizer decreases down the resistance generated by intermolecular forces of polymeric macromolecules[29]. The total volume was made up to 20 mL with distilled water and uniform dispersion was casted on previously prepared backing membrane and dried for 24 hrs. at 37°C.

#### 2.2.3 Statistical optimization of formulated transdermal patch

A four-factor experimental response surface model of four-levelled Box-Behnken design (Design Expert, Version 12, Stat-Ease Inc., Minneapolis, MN) was employed to get the optimized combination of the polymers, solvent and plasticizer (independent variable) to attain the highest yielding percentage drug release (dependent variable). Box-Behnken design has been seen fit to inspect the quadratic response attributes as well as for generating first order polynomial model system[30]. Henceforth, it successfully facilitated the optimization process of a limited numbers of experimental runs. Variables distribution is illustrated in Table 1 and a quadratic design model has been developed via quadratic process order on the basis of experimental data sets as mentioned below and represented in the form of a mathematical expression;

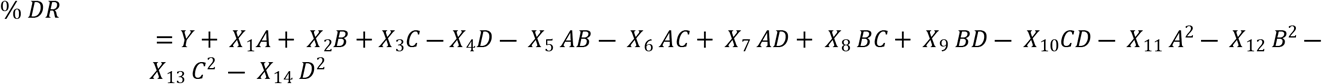

where, percentage drug release has been calculated as the response of the independent variables of the model. Y (86.30) is the intercept whereas X_i_ = (i = 1-13) are the regression coefficients of the experimental model. Moreover the independent variables are denoted as A, B, C and D as mentioned in Table 1 and A^2^, B^2^, C^2^, D^2^ which represents the quadratic terms in the expression[31].

**Table 1:** Box-Behnken variable distribution for experimental design.

| Independent Variable | Levels |  |  |
| --- | --- | --- | --- |
|  | -1 | 0 | 1 |
| A = Polymer 1<br>HPMC concentration (w/v) | 1 | 1.5 | 2 |
| B = Polymer 2<br>PVP concentration (w/v) | 1 | 1.5 | 2 |
| C = Solvent<br>Distilled water (ml) | 10 | 12.5 | 15 |
| D = Plasticizer<br>PEG 400 (w/v) | 0.5 | 0.75 | 1 |
| Dependent Variable | Constraint |  |  |
| DR = Drug Release (percentage) | Maximum |  |  |

Abbreviations: PVP (Polyvinyl pyrrolidone), HPMC (Hydroxypropyl methyl cellulose), PEG 400 (Polyethylene glycol 400).

### 2.3 Characterization of the transdermal patches

#### 2.3.1 Physicochemical parameters of GBP-TDP

The physical attribute plays a vital role in designing an efficient transdermal film and to achieve such results, uniform thickness is desired which was evaluated by measuring thickness at three separate sites employing High-quality Mitutoyo Digimatic micrometer. It is also desirable to remove or attain minimum weight variation, thus, it is analyzed by randomly selecting the five patches and was weighed precisely and calculates the mean average weight to ensure quality observations[32]. Furthermore, the mechanical properties like tensile strength along with elongation of prepared polymeric patches were measured by Tensiometer and the strength of the patches with respect to several polymers and its equilibrated flexibility is determined by number of folds needed to break polymeric patch. Similarly, the cohesive strength exhibited by polymer that determines via evaluating time duration to the pull of adhesive tape and longer the time taken, greater the shear strength [33]. Lastly, the *p*H of the patch was measured wherein, GBP-TDP were kept adjacently in a glass tube with 0.5 mL double distilled water for an hour which further guides patches to swell up[34]. In the wake of such observations glass electrodes were employed which were brought to the close proximity to the patch’s surface to get the *p*H readings however, prior to this process 1 min. of equilibrium period was also given to ensure precise analysis [35].

#### 2.3.2 Percentage Moisture Uptake

Analysis of moisture intake for optimized transdermal patches extends ahead after being kept in CaCl_2_ induced desiccator for 24 hrs. followed by weighing it again and exposing it first to magnesium chloride (MgCl_2_) and then to the solution of sodium nitrite (NaNO_2_) and potassium sulphate (K_2_SO_4_) to stem the presence of relative humidity of 30%, 65% and 97%. Similarly, as discussed earlier these patches’ weight were assessed at regular gap of 48 hrs. until the weight of the patch becomes constant and the calculation of percentage moisture uptake was evaluated using the below mentioned mathematical expression[36]:

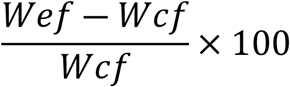

Where, Wef and Wcf are the weights of exposed and conditioned film respectively.

#### 2.3.3 Scanning Electron Microscope

Scanning electron microscope (SEM) was performed to examine and obtain surface-sensitive topographic information and morphology of GBP-TDP for neuropathic pain relief adsorption and effect on the dermal cells. A sample was mounted on High resolution (HR-SEM) support, and then gold coating using a fine coat ion sputter to render the sample electrically conductive in nature, this was further observed optically under SEM (Zeiss EVO, Amity, India) at 100x and 1000x magnification at 20 kV to observe the distribution of GBP on the skin after application of Transdermal patch[37].

#### 2.3.4 Fourier-transform infrared spectroscopy

Fourier-transform infrared spectroscopy (FTIR) was employed to analyse drug-polymer interaction studies for pure drug GBP, physical mixture of PVP and HPMC polymers and transdermal patches using attenuated total reflection (ATR-FTIR) spectroscopy (IR-810. JASCO, Tokyo) at Punjab University, Chandigarh, Punjab, India. Prior analysing sample, the surface of ATR-FTIR was wiped with double distilled water followed by isopropyl alcohol (IPA) and excess of water was gently dried using kim wipes. The transdermal patch was cut in dimensions of 2 cm x 2 cm, and was placed side down onto ATR crystal. For a comprehensive understanding, the test samples were scanned in the range of 400-4000 cm^−1^ [38].

### 2.4 *Ex vivo* permeability kinetics

*Ex vivo* permeation of formulated GBP-TDP was carried out on the goat epithelial membrane. Fresh epithelial tissues were carefully removed from the goat skin obtained from the local slaughterhouse. Defattening and cleaning of epithelial layer was done with PEG 400. The receptor compartment of Franz diffusion cell[39] was filled with PBS (*p*H 5.5) and thin layer of epithelial membrane along with GBP-TDP was mounted on the cell between donor and receptor compartment. The temperature was maintained at 37°C and at predetermined time points, 1 mL samples were withdrawn from the receptor compartment, replacing with equal volume of PBS after each sampling. The samples withdrawn were filtered and used for analysis. The amount of drug permeated was determined using HPLC analysis[40].

The cumulative drug release (CDR %) was estimated by the following expression:

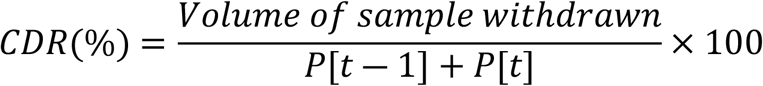

Where, P[t] refers to the percentage release at time [t] and P [t-1] is the percentage release at time [t-1].

## 3. RESULTS AND DISCUSSION

### 3.1 *In silico* validation by targeting hBCATc in the neuropathic pains

#### 3.1.1 Homology Modeling

The 3-D structure of the unknown protein or the target sequence was constructed by aligning a target protein sequence with the template sequence. It was seen that the target with a query coverage of 99% and similarity of 99.74% was obtained having the PDB ID: 2COJ. Later, this target was aligned and modeled with the template for finding out the best model and then validated through Ramachandran Plot showing the most favoured and unfavoured regions and residues of the amino acids. During this iterative process of model building, all the models build are compared and aligned by the program MODELLER and later these models were assessed by a number of criteria depending upon GA341 score and DOPE score. Therefore, for a target sequence to be the best model, the GA341 score equal or near to 1 and the lowest DOPE score is considered[41]. Now this model is regarded as the target for *in silico* docking studies (Figure 1). This target is later validated with the Ramachandran Plot which helps to find out the favoured residues of the amino acid sequence which participate in the docking studies. The Ramachandran plot distinctly represents the phi-psi torsion angles for all residues in the structure and gives the data showing the most favoured and non-favoured regions of the modeled protein. Amino-acid residues are separately identified by the blue triangles scattered all over the four quadrants showing the residues forming the α-helix and β-sheets. As it is clear from the PROCHECK statistics of Ramachandran Plot (Table 1), a good quality model is expected to have over 90% in the most favoured regions [A, B, L] and the observed statistics show that the model has 92% residues in the favoured region possessing 303 residues out of 386 in favoured region. Therefore, on validating through statistics of Ramachandran plot (Figure 2) this model can be used as best model and used as target in docking studies [42].

**Figure 1:**
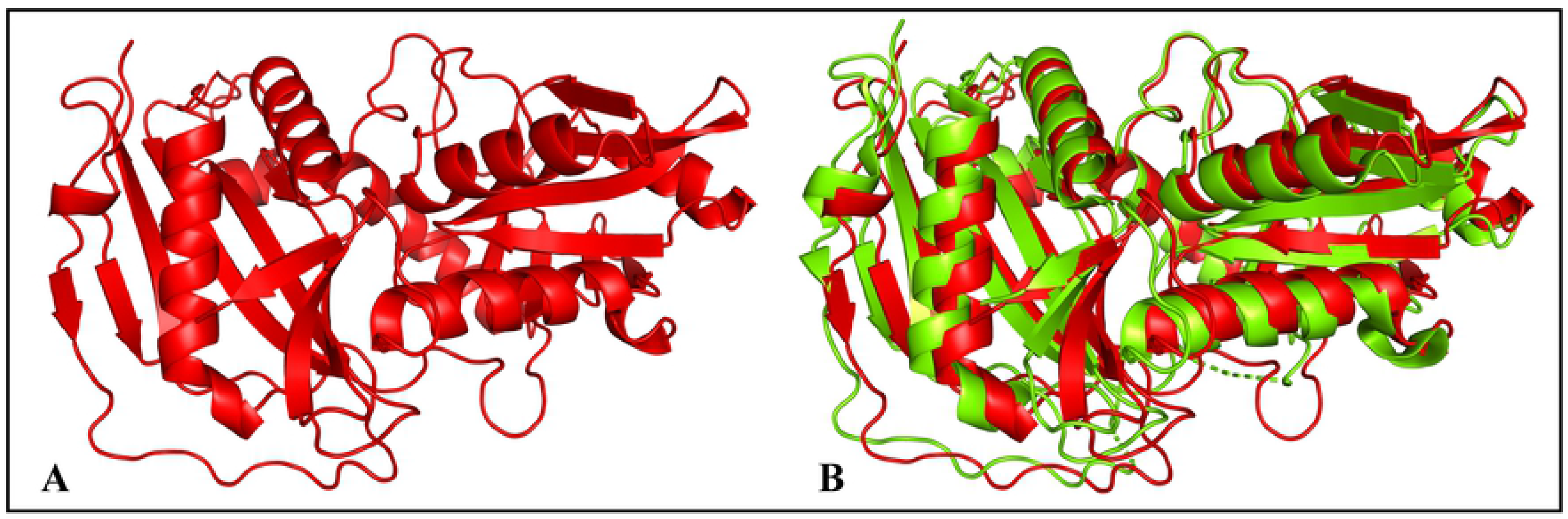
Illustration depicting the modelled protein (A) template protein (2COJ) (B) template protein superimposed to query sequence (BCAT) showing 99% similarity.

**Table 1:** Ramachandran plot statistics

| Parameter | No. of residues | Percentage |
| --- | --- | --- |
| Most favoured regions [A,B,L] | 303 | 92.1% |
| Additional allowed regions [a,b,l,p] | 25 | 7.6% |
| Generously allowed regions [∼a, ∼b, ∼l, ∼p] | 0 | 0.0% |
| Disallowed regions [XX] | 1 | 0.3% |
| Non-glycine and non-proline residues | 329 | 100% |
| End-residues (excl. Gly and Pro) | 2 |  |
| Glycine residues | 33 |  |
| Proline residues | 22 |  |
| Total number of residues | 386 |  |

**Figure 2:**
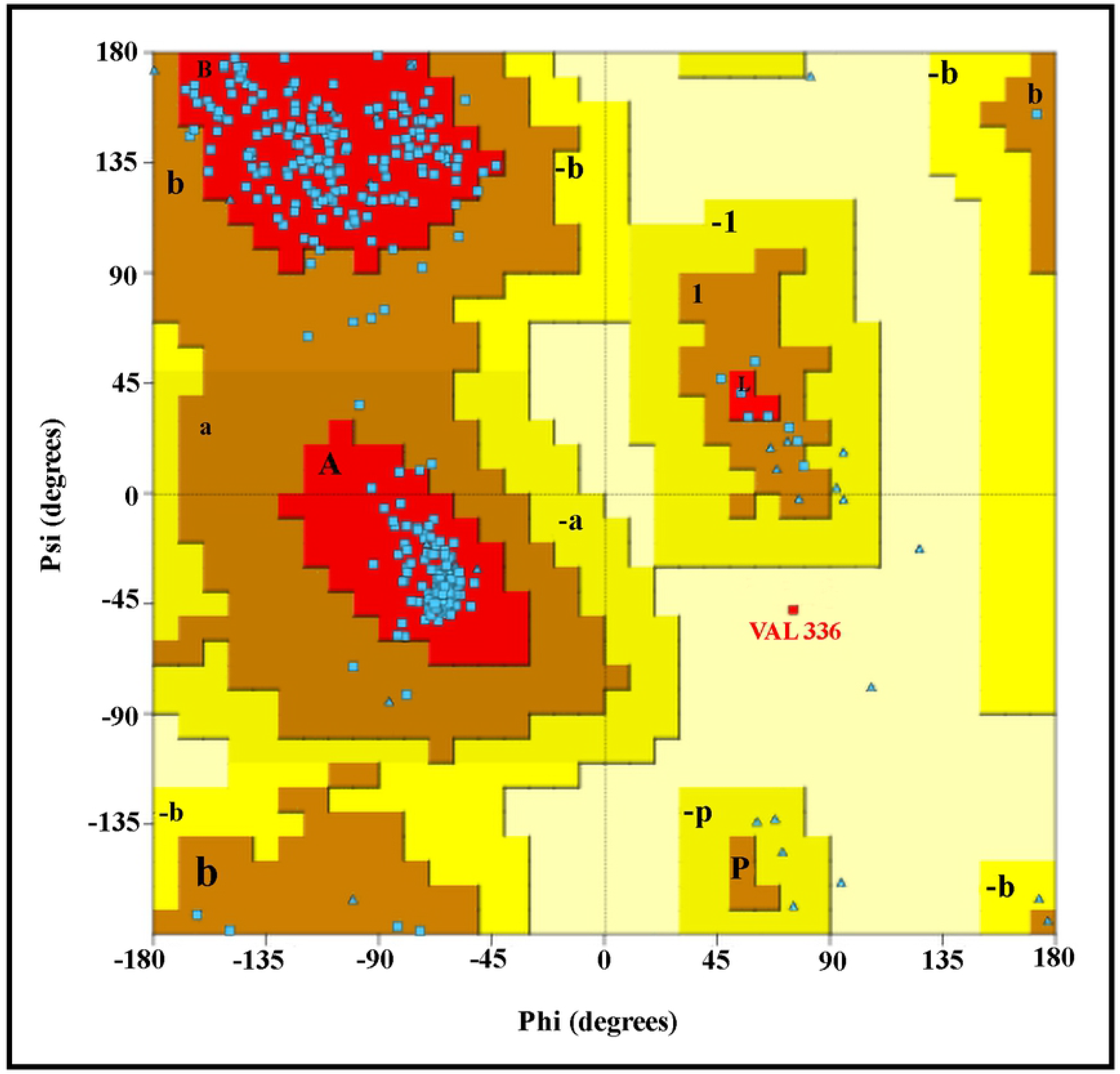
Graph presenting the Ramachandran Plot statistics

#### 3.1.2 Molecular Docking using AutoDock Vina 1.5.6

A total of nine docking stimulations were obtained as results of docking studies of modeled target and the ligand GBP using AutoDock Vina 1.5.6 [43]. The specific binding affinity (in terms of kcal/mol) was used to determine the best docking pose amongst the nine possible docking poses and the one with lowest free energy was considered to be the best conformation of ligand having the highest potential binding affinity toward the active site of the modeled protein. In this case, target ligand complex demonstrated that GBP has highest affinity towards target PDB id: 2COJ with highest specific binding affinity of - 5.3 kcal/mol as shown in Table 2. To have a better visualization of the results and analyze the molecular interactions between the ligand and the modeled target, many computational tools such as Pymol version 2.4, Ligplus version 2.1, UCSF Chimera 1.14 and web based online visualization tools from protein plus website hosted by university of Hamburg (https://proteins.plus/2doo) were used[44]. The target interacting residues at its active site exhibited two distinct types of interaction encompassing hydrogen bonding [Ser157], [Gly191], [Cys335] and Hydrophobic interactions [Thr370], [Val336], [Leu366], [Ser196], [Pro192], [Lys99], [Ala334], [Phe101], [Gln373] as shown in Figure 3. In depth analysis of the molecular interaction, it was observed that the N-atom at C 9 position of ligand with [Ser157] and [Gly191] residues of target formed the salt bridges maintaining a bond length of 3.05 Å and 3.15 Å respectively, screening the hydrogen bond as well as an electrostatic interaction which stabilizes and balances the bond leading to a strong interaction. The second strongest interaction between O2 at C8 position of ligand and O of target residue [Cys335] was identified to be 2.81 Å. Therefore, the present study revealed that the interaction between 2COJ and GBP showed the best docking score by forming strong bonds and regulating their response in tissues when administered.

**Table 2:** Binding affinity of different dock poses.

| Mode | Affinity<br>(kcal/mol) | Distance from best mode |  |
| --- | --- | --- | --- |
|  |  | R.M.S.D l.b | R.M.S.D u.b |
| 1 | -5.3 | 0.000 | 0.000 |
| 2 | -5.2 | 13.105 | 13.893 |
| 3 | -5.1 | 2.479 | 4.293 |
| 4 | -5.1 | 1.644 | 2.676 |
| 5 | -5.0 | 11.926 | 13.015 |
| 6 | -5.0 | 2.093 | 2.834 |
| 7 | -4.9 | 2.922 | 4.228 |
| 8 | -4.9 | 3.164 | 4.403 |
| 9 | -4.9 | 12.019 | 13.118 |

**Figure 3:**
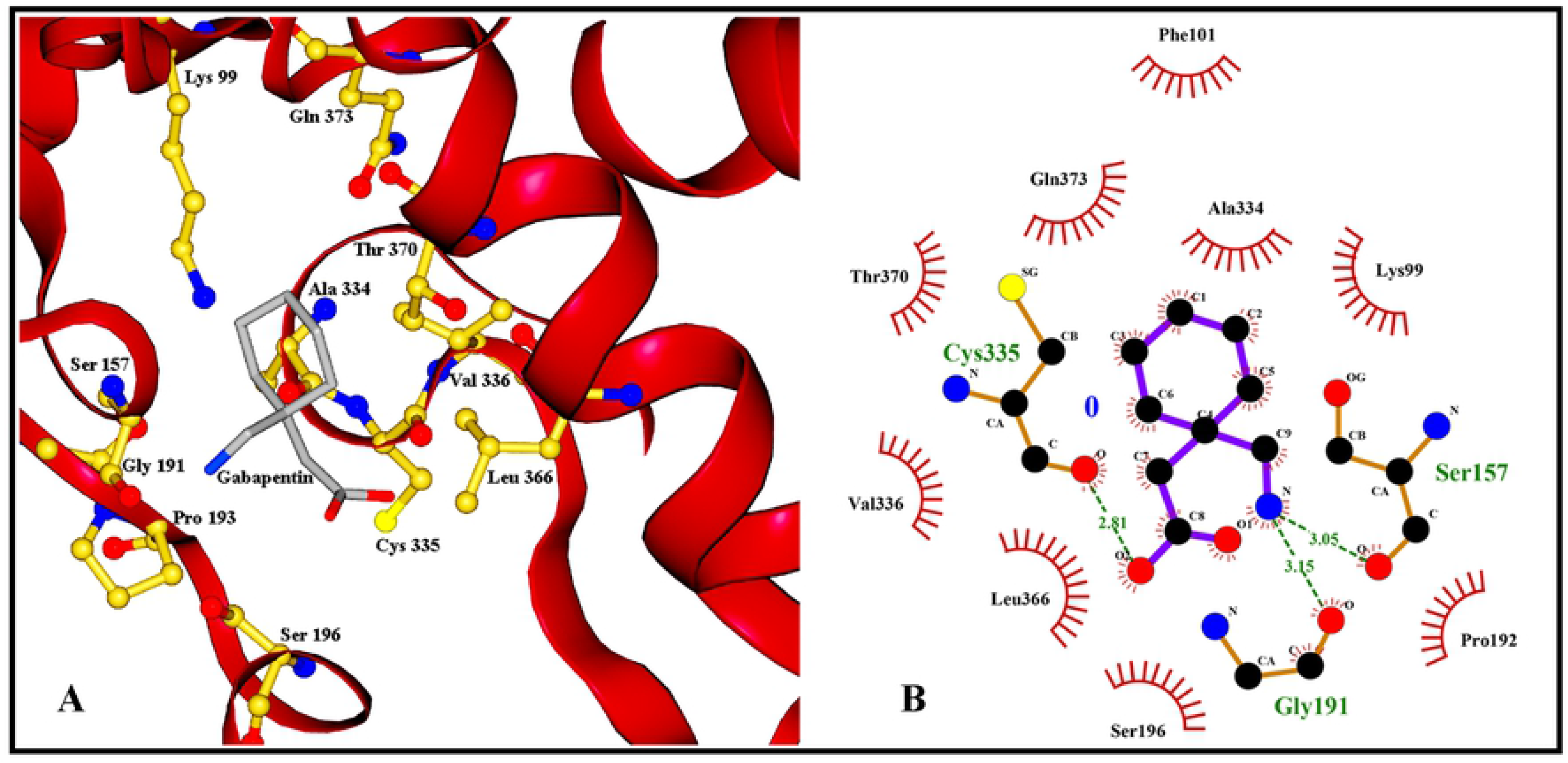
Docking results of 2COJ (human branched chain aminotransferases) and their interacting residues with Gabapentin

#### 3.1.3 ADMET

In this profiling, we retrieved that GBP inhibits the release of excitatory neurotransmitters and possesses 12 heavy atoms, 3 hydrogen bond donors and 2 acceptors in its structure[45]. Further, its bioavailability radar (Figure 4) shows six physiochemical properties (size, lipophilicity, polarity, saturation, solubility and flexibility) and the area in pink colour depicts the descriptors that falls in entirely in plot confirming the drug-like properties of GBP (Table 3). Similarly, the lipophilicity of GBP (log P_o/w_) from different predictive models of iLOGP and XLOGP3 is 1.45 and −1.41 respectively, confirming that it has good absorption and permeation in the tissues for drug delivery through transdermal patches. Also, the consensus log P_o/w_ shows the arithmetic mean of all the predicted values from all other descriptors as 0.62, further assuring the accuracy of the result. Thereafter, the aqueous solubility (log S) or ESOL model, implemented to find out the general solubility equation, gives R^2^ value of 0.18 suggesting that the drug is highly soluble and would have enhanced absorption and distribution characteristics. Lastly, GBP also reflected its suitability for drug like properties as per the Lipinski’s rule [46].

**Table 3:** ADMET profiling.

| SMILES | NNC1CCCC(C1)CC(=O)O |
| --- | --- |
| **Physicochemical Properties** |  |
| Formula | C<sub>9</sub>H<sub>17</sub>NO<sub>2</sub> |
| Molecular weight | 171.24 g/mol. |
| No. heavy atoms | 12 |
| No. H-bond acceptors | 3 |
| No. H-bond donors | 2 |
| Molar refractivity | 47.74 |
| TPSA | 63.32 Å<sup>2</sup> |
| **Lipophilicity** |  |
| Log P<sub>o/w</sub>(iLOGP) | 1.45 |
| Log P<sub>o/w</sub>(XLOGP3) | -1.41 |
| Consensus Log P<sub>o/w</sub> | 0.62 |
| **Water Solubility** |  |
| Log S(ESOL) | 0.18 |
| Solubility | 2.62e+02 mg/ml;<br>1.53e+00 mol./l |
| Class | Highly soluble |
| **Pharmacokinetics** |  |
| GI absorption | High |
| BBB permeant | Yes |
| P-gp substrate | No |
| CYP1A2 inhibitor | No |
| CYP2C19 inhibitor | No |
| CYP2C9 inhibitor | No |
| CYP2D6 inhibitor | No |
| CYP3A4 inhibitor | No |
| Log K<sub>p</sub> (skin permeation) | No |
| **Drug-likeness** |  |
| Lipinski | Yes |
| Bioavailability score | 0.55 |

**Figure 4:**
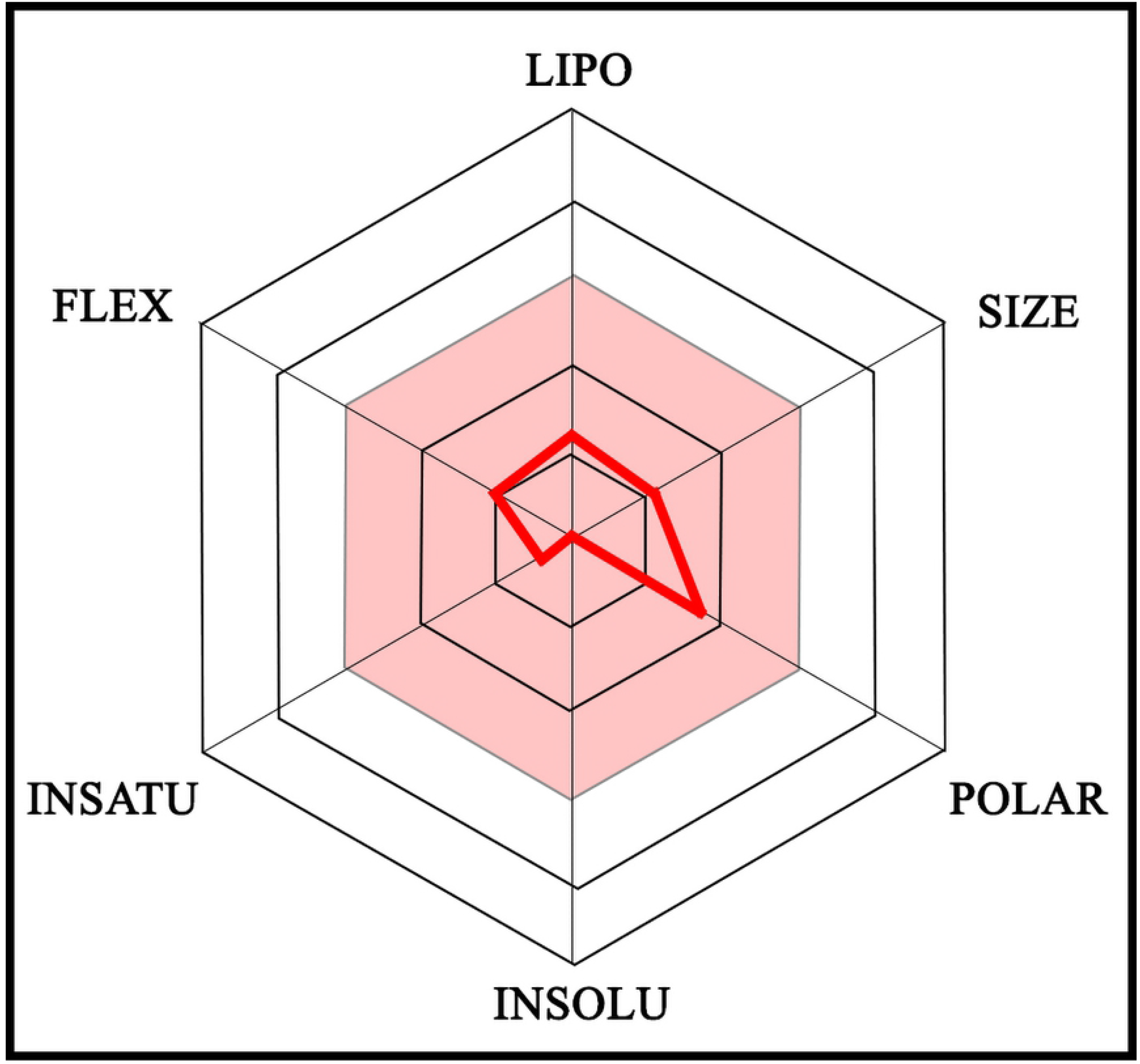
Oral bioavailability radars showing: LIPO (lipophilicity): −0.7 < XLOGP3 < +5.0; SIZE: 150g/mol < MV < 500g/mol; POLAR (polarity): 20Å^2^ < TPSA < 130Å^2^; INSOLU(insolubility): 0< Log S (ESOL) <6; INSATU(insaturation): 0.25 < Fraction Csp3 < 1 and FLEX(flexibility): 0 < Num. rotatable bonds < 9.

### 3.2 Preparation of GBP-TDP

The backing membrane was prepared using the solvent evaporation method and the membrane prepared by dissolving 4% (w/v) PVA was found to be most suitable, transparent and flexible. After the fixation of backing membrane, the patches composed of PVP (2%) and HPMC (1%) with PEG 400 (0.5%) as plasticizer and DMSO as permeation enhancer were further casted and analysed for their various physicochemical parameters and statistical optimisation as shown in Figure 5.

**Figure 5:**
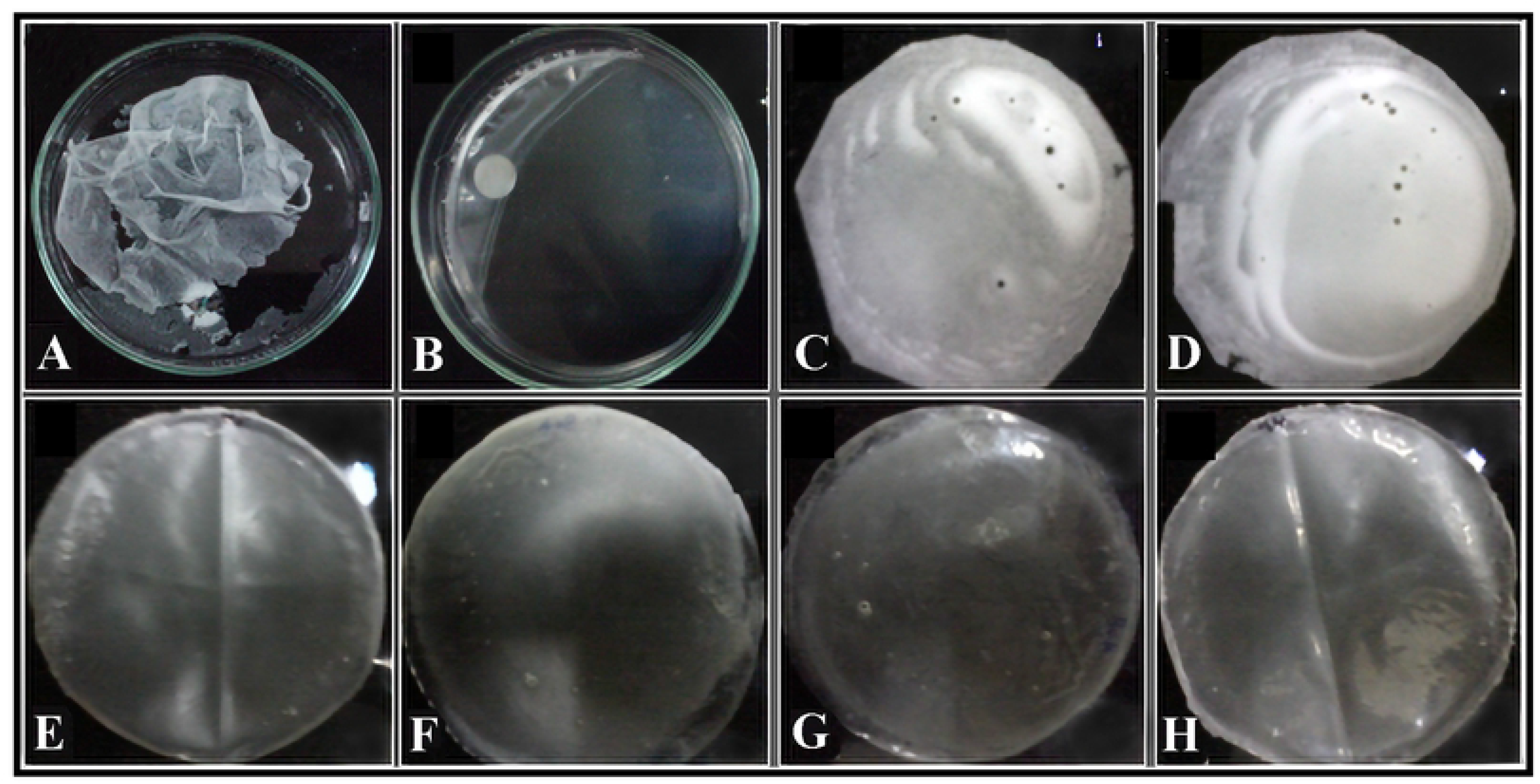
The transdermal patch casting images exhibiting GBP – TDP with varied types of polymer concentrations wherein PVP:HPMC combination are: (A) 0.5:0.5, (B) 0.5:1, (C) 1:0.5, (D) 1:1, (E) 1.5:1, (F) 2:1, (G) 1:1.5, (H) 1:2.

#### 3.2.1 Statistical optimization of transdermal patches

Four-levelled Box-Behnken model conducted 29.00 experimental randomized runs with varying combinations of independent variables and the p-value 0.0455. This run also fulfils the requirement of the minimum number runs for generating the RSM model as well[47]. Designed system was observed to producing highly significant data with R^2^ value of 0.3226 and F-value of 2.86 as shown in Table 4. Moreover, on the analysis of the mathematical expression of the polynomial model with quadratic process order we came to infer that ratios of independent factors B and D have seemed to be directly proportional, however factors A and C influence the release of the drug indirectly. Thus, a cumulative effect of the contributing independent variables on the percentage drug release can be explained by the mathematical expression:

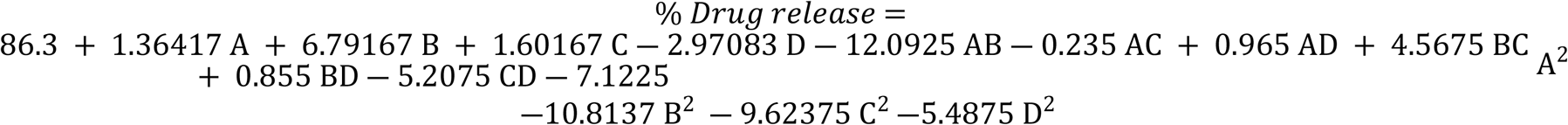

**Table 4:** Analysis of variance (ANOVA) for the percentage drug release.

| Source | Sum of squares | df | Mean square | F-value | p-value | R<sup>2</sup> value |  |
| --- | --- | --- | --- | --- | --- | --- | --- |
| **Model** | 941.89 | 4 | 235.47 | **2.86** | **0.0455** | **0.3226** | **significant** |
| A-HPMC concentration | 0.5334 | 1 | 0.5334 | 0.0065 | 0.9365 |  |  |
| B-PVP concentration | 803.77 | 1 | 803.77 | 9.75 | 0.0046 |  |  |
| C-Plasticizer PEG 400 | 30.78 | 1 | 30.78 | 0.3735 | 0.5468 |  |  |
| D-Solvent distilled water | 106.80 | 1 | 106.80 | 1.30 | 0.2662 |  |  |
| Lack of Fit | 1977.88 | 20 | 98.89 |  |  |  |  |
| Pure Error | 0.000 | 4 | 0.000 |  |  |  |  |

Moreover, the significance of the data sets as mentioned in Table 2 further suggests that the selected optimized combination of 1% HPMC, 2% PVP, 0.5% plasticizer and 10 mL of solvent (in the 14^th^ run) was observed to yielding the highest drug release of 93.74% from the synthesized transdermal film. Normal distribution and linear regression of the designed model was ensured by running a normal residual plot (Figure 6 (A)). Desirability of a data is viewed as an objective function which simply ranges from 0–1 at the projected or decided goal for the study. The 3-D plot here clearly establishes the numerical optimization of our dataset which came out to be 1 (Figure 6 (B)); thus the suitability and significance of the response with respect to the dataset is high as there aren’t any deviations in the response for a distinct combination of independent variables. Furthermore, the response surface methodology (RSM) plots, reinforces the significance of the data generated in ANOVA studies (Figure 6 (c)).

**Figure 6:**
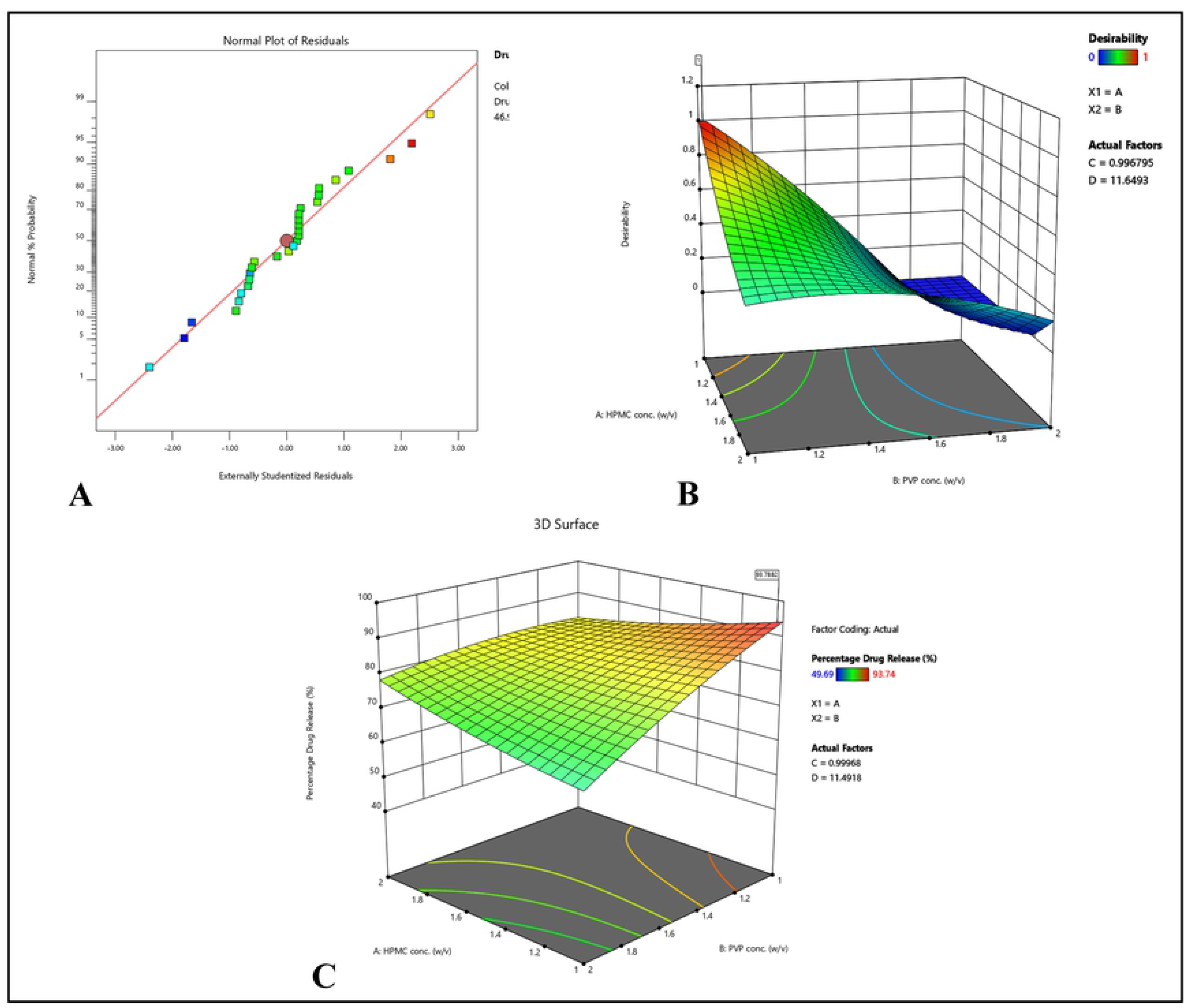
Images representing the polymer interaction plots where image (a) shows predicted versus experimental graph of encapsulation efficiency; (b) and (c) represents three-dimensional surface interaction plots showing various interactions with respect to encapsulation efficiency and desirability ratios between the polymers to attain ionic gelation matrix

### 3.3 Characterization of transdermal patch

#### 3..3.1 Physicochemical properties

GBP-TDP was evaluated for a variety of physico-chemical analysis which helped to understand the physical state of the drug delivery system. The physico-chemical studies like thickness, weight variation, tensile strength, folding endurance, and shear adhesion of patch 9 was calculated and they were found to be uniform, flexible, transparent and smooth. The weight of the formulated patch was 19.2 ± 0.12 mg and the thickness was 0.0234 ± 0.0002 mm as shown in Table 5. Low standard deviation between the patches ensured uniformity of the patches and low variability in the reproducibility of the procedure followed for the preparation of patches. The folding endurance value was found to be 60 ± 2.44. The results of folding endurance value indicate that the patch was found to be satisfactory which indicated that the patch prepared using PEG 400 and the polymer concentration of 4% (PVA) had an optimum flexibility and were not brittle. In another study, where Prajapati et al, prepared the transdermal patch of Repaglinide, the folding endurance ranged between 20-82 which further validates our result[48]. The tensile strength and shear adhesion of the patch 9 was measured to be 2.3 ± 0.23kg/mm^2^ and 0.5 kN/m respectively which demonstrated the durability and stability to stress and endurance to external forces (Table 5). This also suggests the strong cohesive force due high degree of uniform crosslinking of polymers and optimal composition and flexibility of patch. Lastly, the neutral *p*H of the patch makes it ideal to be applied on the skin for the delivery of GBP.

**Table 5:** Physico-chemical parameters of the optimized patch.

| Parameters | Thickness (mm) | Folding Endurance | Weight (mg) | Surface pH | Shear Adhesion (hr.) | Tensile Strength (kg/mm<sup>2</sup>) |
| --- | --- | --- | --- | --- | --- | --- |
| Patch P9 | 0.0234 ± 0.0002 | 300 ± 2.44 | 1.92 ± 0.12 | 6.8 ± 1.23 | 0.5 ± 0.003 kN/m | 2.3 ± 0.23 |

#### 3.3.2 Moisture Uptake study

The moisture content of the optimized patch was done to assess the integrity of the films at different humid conditions (33%, 65% and 97%) after 24, 48 and 72 hrs. The prepared patches exhibited minimal water absorption rate as shown in Figure 7 which ensures stability and prevents microbial contamination. The lower moisture content even at higher relative humidity helps the patches to become completely dried and brittle film [49].

**Figure 7:**
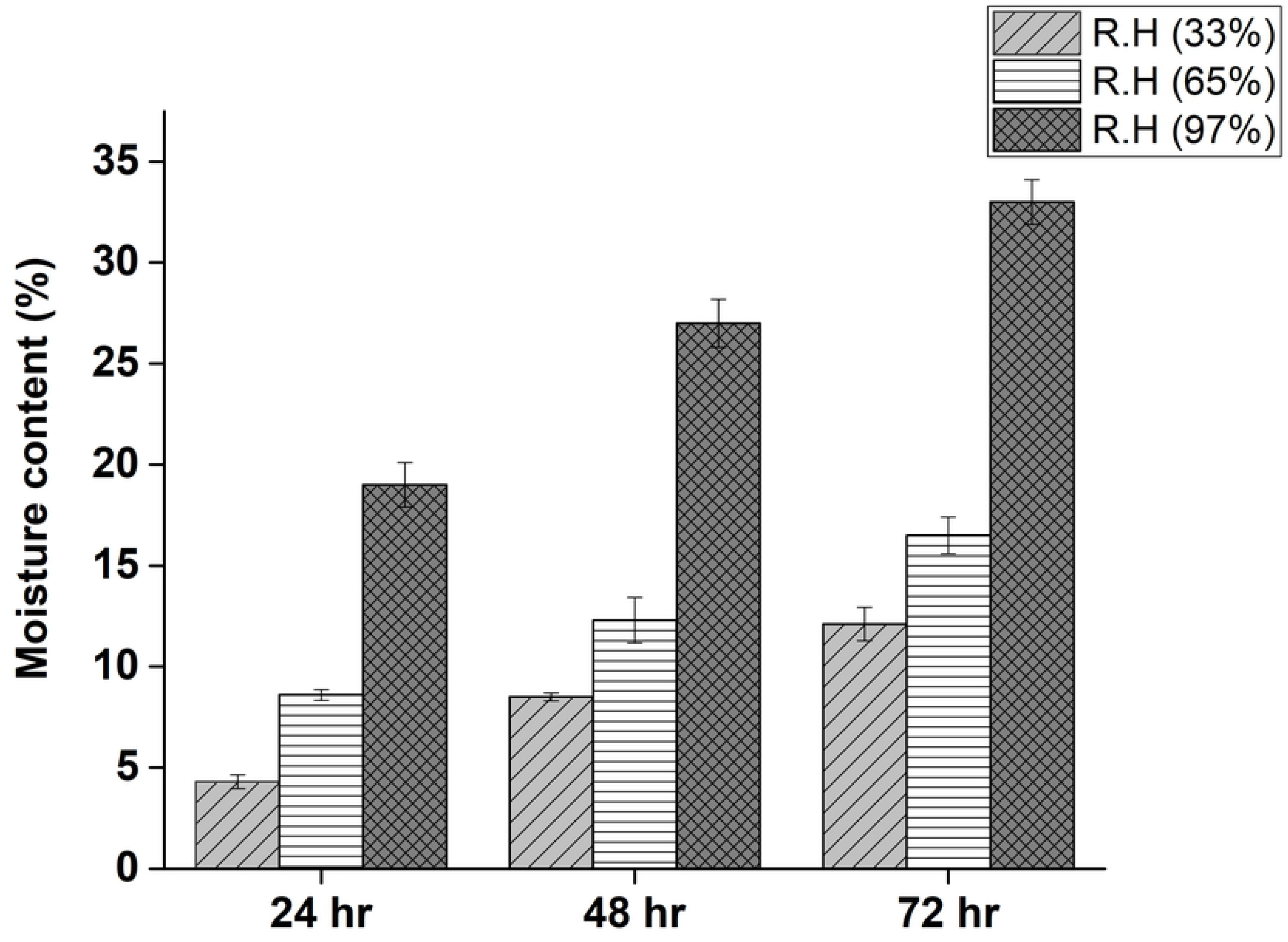
Water uptake study by exposing the patches to 33%, 65% and 97% relative humidity.

#### 3.3.3 Scanning Electron Microscope (SEM)

In order to optically visualize the effects of novel GBP-TDP, we employed high resolution Scanning Microscope of dermal cells treated with the transdermal patch. As shown in Figure 8, we can clearly observe there was no deformation, damage or distortion of the patch layer which are consistent with the result where the permeation occurs without damaging the epidermal layer. Figure 8 (A and B) demonstrates the uniform distribution and presence of drug in transdermal formulation and easy permeation was achieved.

**Figure 8:**
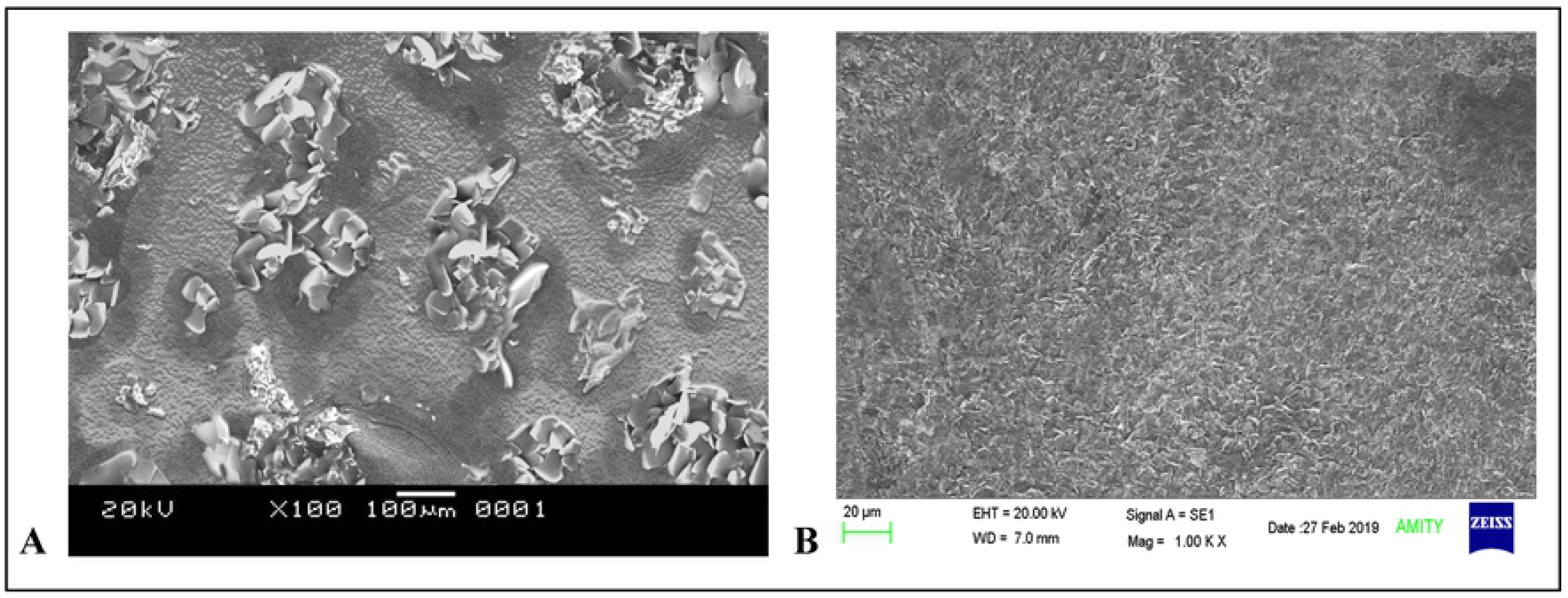
High resolution-scanning electron micrograph of GBP-TDP at the magnification of (a) 100x (left); (b) 1000x (right).

#### 3.3.4 Fourier-transform infrared spectroscopy analysis

GBP is a cyclohexane and consists of COOH, NH_2_ group. The O-H stretch and C=O stretch of -COOH is present at 2923 cm^−1^, 1740 cm^−1^ in the range of 3000-2500 cm^−1^ respectively. The bands between 1650–1400 cm^−1^ are assigned to carbon vibrations whereas the C-C stretching vibrations interact with C-H in plane bending vibrations is observed in the range of 1300-1000 cm^−1^ (Figure 9). The amine group of GBP molecule is involved in the formation of covalent bond with miscible blend of PVC/HPMC polymer. In this, −NH stretch situated at 3279 cm^−1^ in the range of 3100-3500 cm^−1^ is of great importance, this amide functional group combines with the feature of >C=O and >N-C groups at 1633 cm^−1^ and 1309 cm^−1^ of PVP polymer. Therefore, amide bond shows strong somewhat broad bend at left hand side of spectrum. On the other hand, the characteristic peaks of HPMC are at 3050-3750 cm^−1^ which superpositions amide stretch, signifying the characteristic peak to follow the same path. A broad bend at 3279 cm^−1^ denotes overlapping of two types of −OH group present in HPMC polymer-free and self-association −OH group at 3585 cm^−1^ forming intermolecular and intramolecular H-bonding at 3362 cm^−1^ which cannot be observed due to peak broadness and embedment of GBP inside the patch. But, the positioning and intensity of these characteristic peak permit rather encourage to interpret interactions of these prepared polymers with GBP have been fully blended on the surface.

**Figure 9:**
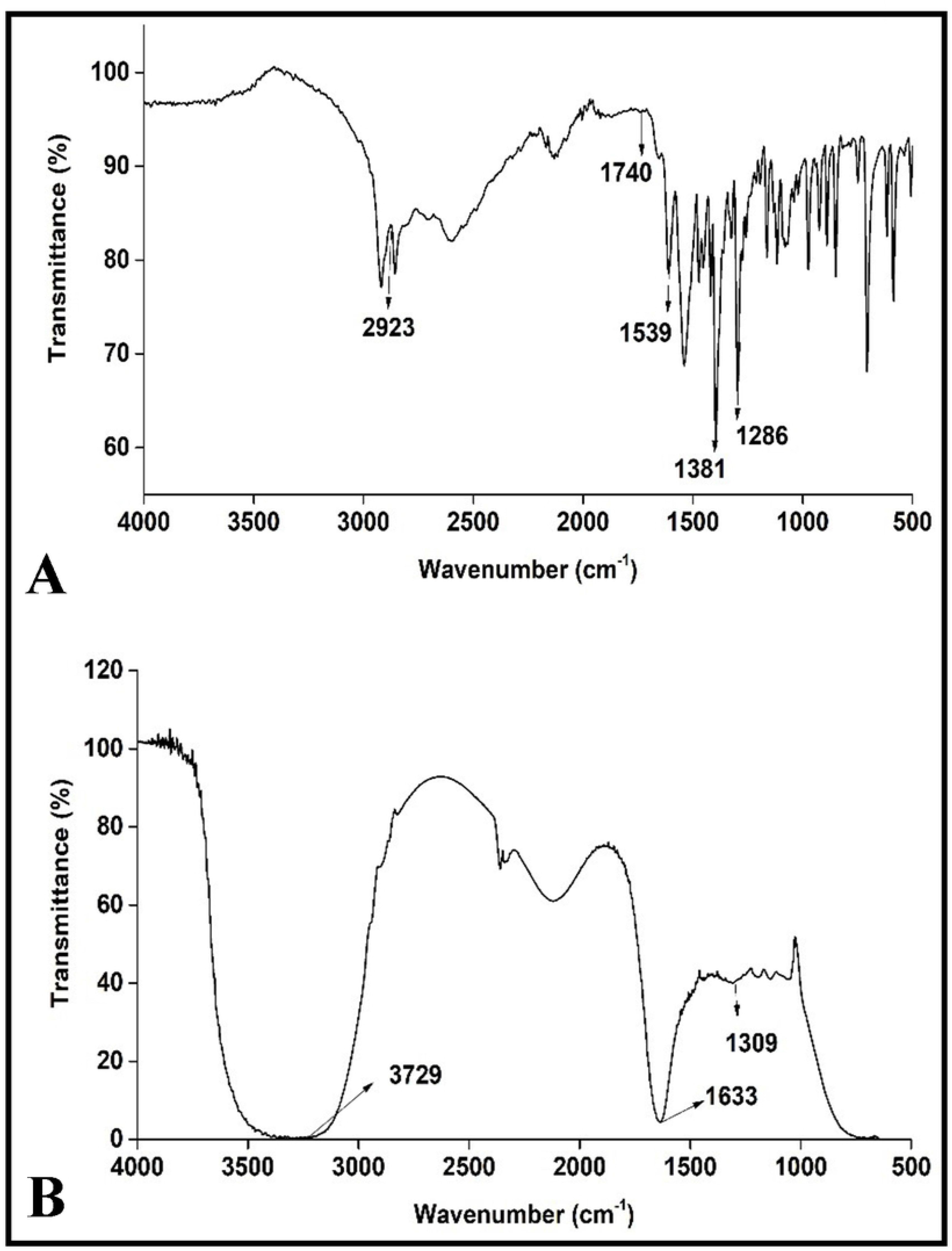
FTIR spectra of (A) GBP and (B) GBP-TDP.

### 3.4 Drug release kinetics study

The *ex vivo* release kinetics of GBP and optimized formulation of GBP-TDP were analyzed using an impermeable membrane. The release kinetics data revealed the cumulative drug release of 92.34 ± 1.43% of GBP from the transdermal patch after 8 hrs. while only 60.87 ± 0.54% was released in case of GBP. The optimized GBP-TDP showed slow release of GBP from the patch because of the matrix formation which followed zero order kinetics (R^2^= 0.9835). The zero order kinetics represents that the drug does not disintegrate and is released slowly from the biological membranes as shown in Table 6. Furthermore, there was a sharp decrease at 5^th^ hour with cumulative drug release of 35.71 ± 1.24% in case of GBP as shown in Figure 10. The comparative study of results obtained for the test samples revealed improvement in the release characteristics of GBP from a patch.

**Table 6:** Kinetic models for *ex vivo* release of GBP and GBP-TDP.

| Formulation | Zero order | First order | Higuchi | Hixson | Korsmeyer-Peppas |
| --- | --- | --- | --- | --- | --- |
| GBP | 0.217 | 0.075 | 0.0541 | 0.1284 | 0.1062 |
| GBP-TDP | 0.9835 | 0.8007 | 0.9475 | 0.9419 | 0.9357 |

**Figure 10:**
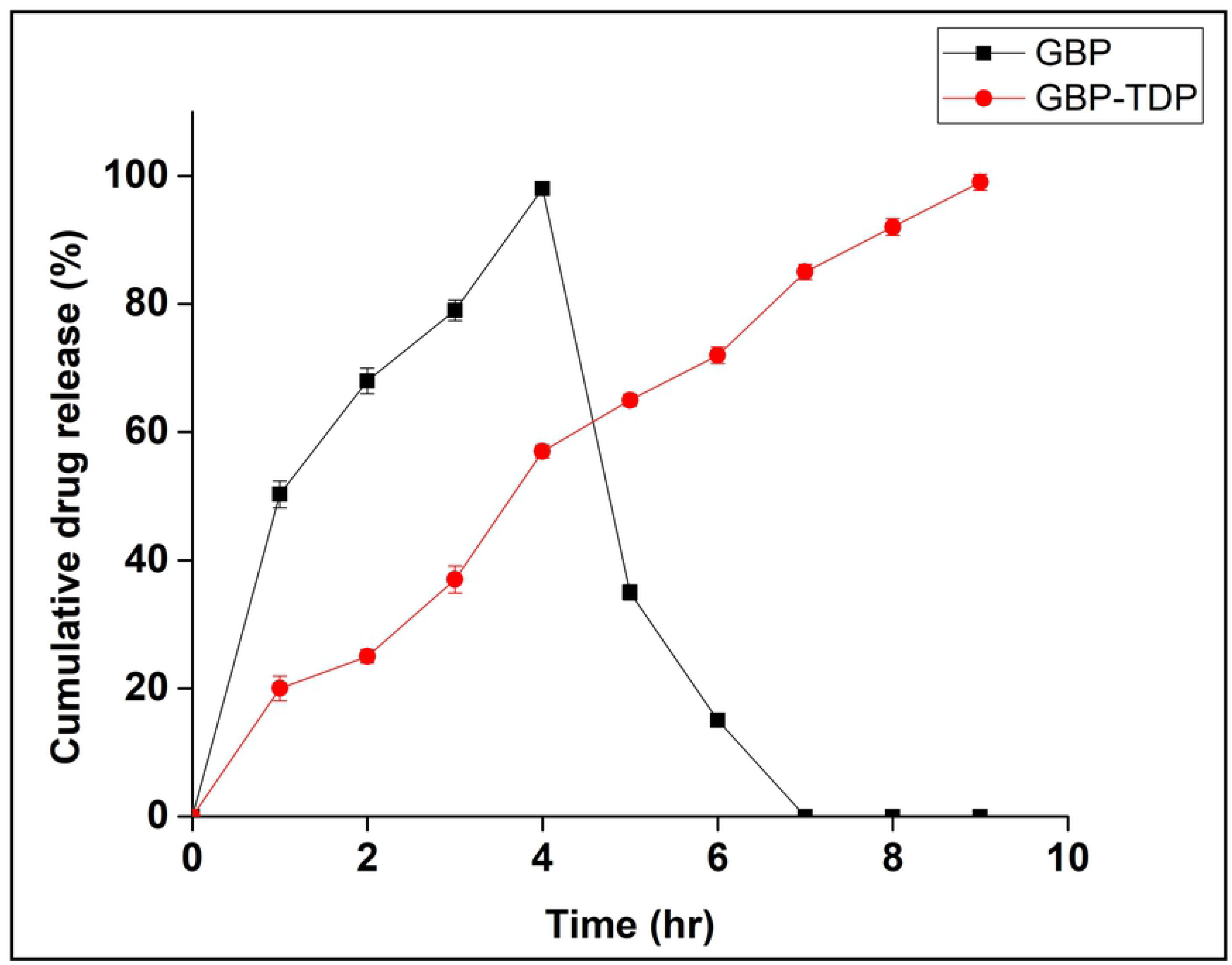
*Ex vivo* release kinetics of GBP and optimized GBP-TDP.

## 4. CONCLUSION

Transdermal drug delivery systems are designed to deliver drugs to dermal layer that permeates deeper into the systemic circulation directly avoiding the hepatic first pass metabolism. In this study, we have repurposed Gabapentin to treat diabetic neuropathy which has been validated by using various bioinformatics tool. We prepared transdermal drug delivery system of Gabapentin using different combinations of polymers (PVP and HPMC), plasticizer (PEG 400) were synthesized using solvent evaporation method. Thereafter, four-levelled Box-Behnken design was employed to get the optimized combination of the polymers, solvent and plasticizer to attain the highest yielding percentage drug release. The optimized formulation consisted of PVP and HPMC at the ratio of 2:1, PEG 400 and PVA (4%). Thereafter, the prepared films were evaluated in terms of physical appearance, uniformity, weight, tensile strength, surface *p*H, moisture content and uptake, the results suggested that the transdermal patches were of excellent quality and were highly reproducible with minimum variability. The transdermal patches were characterized by Scanning electron microscopy and Fourier transform Infrared Spectroscopy which depicts the uniform distribution of patches and the successful attachment of Gabapentin onto the patches. Further, *ex vivo* drug release kinetics of Gabapentin-transdermal patch followed the zero order kinetics and nearly complete release of Gabapentin after 8 hrs. in comparison to burst release of GBP alone. These results show that the transdermal delivery of Gabapentin can have potential applications in diabetic neuropathy and can offer various advantages like improved bioavailability, non-invasive delivery, improved patient compliance, reduced dosing frequency.

## Acknowledgments

The research group is grateful to the Department of Biotechnology, Jaypee Institute of Information Technology, Noida, Uttar Pradesh for providing necessary facilities to execute this work.

